# Three-dimensional Shear Wave Elastography Using a 2D Row Column Addressing (RCA) Array

**DOI:** 10.1101/2021.11.10.467798

**Authors:** Zhijie Dong, Jihun Kim, Chengwu Huang, Matthew R. Lowerison, Shigao Chen, Pengfei Song

**Author notes:** Corresponding Author: Dr. Pengfei Song, Department of Electrical and Computer Engineering, Beckman Institute for Advanced Science and Technology, University of Illinois Urbana-Champaign, 405 N. Mathews Ave. Urbana, IL 61801.

## Abstract

**Objective:** To develop a 3D shear wave elastography (SWE) technique using a 2D row column addressing (RCA) array, with either external vibration or acoustic radiation force (ARF) as the shear wave source.

**Impact Statement:** The proposed method paves the way for clinical translation of 3D-SWE based on the 2D RCA, providing a low-cost and high volume-rate solution that is compatible with existing clinical systems.

**Introduction:** SWE is an established ultrasound imaging modality that provides a direct and quantitative assessment of tissue stiffness, which is significant for a wide range of clinical applications including cancer and liver fibrosis. SWE requires high frame-rate imaging for robust shear wave tracking. Due to the technical challenges associated with high volume-rate imaging in 3D, current SWE techniques are typically confined to 2D. Advancing SWE from 2D to 3D is significant because of the heterogeneous nature of tissue, which demands 3D imaging for accurate and comprehensive evaluation.

**Methods:** A 3D SWE method using a 2D RCA array was developed with a volume-rate up to 2000 Hz. The performance of the proposed method was systematically evaluated on tissue-mimicking elasticity phantoms.

**Results:** 3D shear wave motion induced by either external vibration or ARF was successfully detected with the proposed method. Robust 3D shear wave speed maps were reconstructed for both homogeneous and heterogeneous phantoms with inclusions.

**Conclusion:** The high volume-rate 3D imaging provided by the 2D RCA array provides a robust and practical solution for 3D SWE with a clear pathway for future clinical translation.

## Introduction

Shear wave elastography (SWE) is an established ultrasound imaging modality that found significant impact in a wide range of clinical applications including cancer and liver fibrosis [1–4]. SWE estimates the speed of fast propagating shear waves in tissue, which is directly related to tissue stiffness (i.e., shear modulus) and provides an important biomarker for tissue health [5–11]. Shear waves are typically generated by either using the acoustic radiation force (ARF) from the push beam of an ultrasound transducer [12], or by using an external mechanical vibration coupled to the tissue [13]. Due to the fast propagating speed of shear waves (e.g., 1 – 10 m/s in most soft tissues), a high imaging frame rate (e.g., 1000-2000 Hz) is typically required for robust SWE [14]. While such high imaging speed is now widely available on mainstream ultrasound scanners powered by software beamformers for 2D imaging, a viable 3D imaging solution that supports such high imaging speed (i.e., >1000 Hz volume-rate) remains elusive. As such, current SWE implementations are largely limited to 2D, although 3D SWE has many known advantages over 2D because of the anisotropic tissue mechanical properties and the resulting complex shear wave propagations in 3D [15]. Furthermore, 2D imaging is particularly ill-posed for external vibration-induced shear waves because of the complexities associated with the shear wave source and various boundary conditions [16].

To date, several solutions for 3D SWE have been proposed. One category uses mechanical scanning of 1D transducers (e.g., wobblers) [17] to acquire 2D SWE images (usually ARF-based) at different elevational slices followed by 3D reconstruction. These methods are not true 3D-SWE given that shear wave detection and tracking are still performed in 2D (i.e., within each 2D imaging slice), which does not carry information of tissue viscoelasticity orthogonal to the 2D imaging plane. However, wobbler-based 3D-SWE is commercially available and readily accessible to clinicians worldwide [18].

The other main category of 3D-SWE methods is based on 2D matrix arrays, with either external vibration or ARF as the shear wave source. Although a previous study reported the use of ARF for 3D-SWE on a 32 x 32 element 2D array [19], in general, it is challenging to implement such an approach on 2D arrays with multiplexing [20], as multiplexed 2D arrays cannot sustain the long-duration, high-voltage push pulse for ARF. In addition, ARF-induced shear waves are transient and wideband (e.g., 75 – 500 Hz) [21], which requires a high volume rate imaging for tracking (e.g., 1000 – 2000 Hz). Due to the high channel count and expensive computational cost associated with 3D beamforming, it is challenging to reach such high volume-rate with 3D-SWE.

On the other hand, external vibration-based methods are more feasible for 3D-SWE based on 2D arrays because they do not use ARF and the shear wave signal is typically lower in frequency and narrower in bandwidth. In particular, for methods using continuous vibrations, thanks to the cyclic nature of the resulting shear wave signal, one can use sub-apertures of the transducer to acquire sub-volumes of the shear wave signal separately and then stitch the sub-volumes of shear wave signals together to synthesize the full 3D volume [22, 23]. The cyclic nature of the continuous vibration-induced shear waves can be further exploited to enable 3D-SWE on a conventional ultrasound scanner with a very low imaging volume-rate. For example, Huang *et al*. [24] proposed a sub-Nyquist sampling technique that allows tracking of 3D shear waves with an 88.9 Hz volume rate. However, one common issue with these sub-volume and sub-Nyquist sampling-based methods is that multiple data acquisitions are necessary to synthesize the data for the full 3D volume, which makes these methods susceptible to artifacts from tissue motion.

To address these limitations, here we propose a new 3D SWE method based on the 2D row column addressing (RCA) array. Different from the design of a conventional 2D matrix array (e.g., N × N elements), RCA arrays use orthogonally arranged, bar-shaped elements (e.g., N + N elements) to reduce the fabrication and computational complexities of 3D imaging. Thanks to the low channel count, RCA arrays are compatible with most commercial ultrasound scanners and support ultrafast, 3D volumetric imaging with thousands of Hertz of volume rate [17, 25]. Meanwhile, thanks to the absence of multiplexing, RCA arrays can sustain the push pulses for ARF-based 3D SWE. As such, 2D RCA arrays present an enticing solution for robust 3D-SWE. Although several studies have presented methods for 3D ultrafast blood flow and super-resolution imaging based on RCA [26, 27], to the best of our knowledge there have been no studies reporting the use of RCA for 3D-SWE. In this paper we will first present a study that investigates different plane wave compounding schemes based on the RCA for 3D shear wave tracking, followed by phantom studies that demonstrate 3D-SWE using the RCA for both external vibration- and ARF-induced shear waves.

## Results

### Imaging sequence for 3D shear wave detection

Three compounding plane wave imaging methods were studied based on the RCA array. The first scheme is compounding RC, as shown in Figure 1(a), which uses row elements (R) to transmit plane-waves with different steering angles and column elements (C) to receive (i.e., the RC scheme). Similarly, the CR scheme uses columns (C) to transmit and rows (R) to receive. The compounding RC+CR combines RC and CR and achieves symmetrical PSF (see ‘Imaging sequence for 3D shear wave detection’ in Materials and Methods). The specifications of the sequences are listed in Table I.

**Table I.** Sequence specifications for different transmission schemes (Central frequency =5.2 MHz; FOV = 25.6 × 25.6 × 50 mm^3^).

| Specs | Transmission scheme |  |  |  |  |  |
| --- | --- | --- | --- | --- | --- | --- |
|  | 400 Hz |  |  | 2000 Hz |  |  |
|  | Compounding RC | Compounding CR | Compounding RC+CR | Compounding RC | Compounding CR | Compounding RC+CR |
| Number of transmissions | 31 | 31 | 30 | 7 | 7 | 6 |
| Angular pitch [°] | 1 | 1 | 1 | 1 | 1 | 1 |
| Maximum angular aperture [°] | 15 | 15 | 7 | 3 | 3 | 1 |
| Transmit focusing | y-direction | x-direction | x- and y-direction | y-direction | x-direction | x- and y-direction |
| Receive focusing | x-direction | y-direction | x- and y-direction | x-direction | y-direction | x- and y-direction |
| Pulse repetition frequency, PRF [kHz] | 12.4 kHz | 12.4 kHz | 12 kHz | 14 kHz | 14 kHz | 12 kHz |

**Figure 1:**
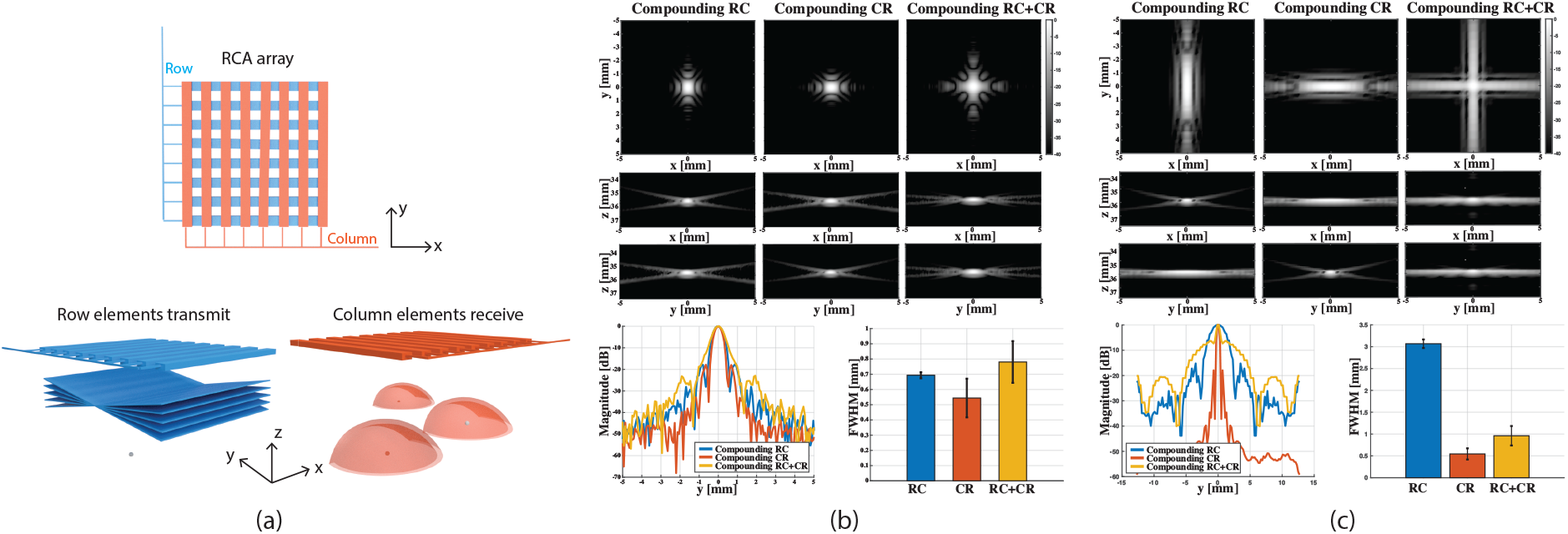
Illustration of the transmission scheme and simulated point spread functions (PSFs). (a) Illustration of the compounding RC scheme of the RCA array. The row elements (distributed along the y-direction) transmit steered plane waves into the medium, and the column elements (distributed along the x-direction) receive the backscattered signals (spherical waves emitted by three example scatterers). Transmit focusing is achieved in the y-z plane and receive dynamic focusing is achieved in the x-z plane. (b) Simulated PSFs of different transmission schemes (compounding RC, compounding CR, and compounding RC+CR) at the volume rate of 400 Hz, as listed in Table I. Profiles of the PSFs and the averaged FWHMs along the y-direction for different transmission schemes are shown in the bottom. (c) Simulated PSFs of different transmission schemes at the volume rate of 2000 Hz, as listed in Table I.

Figure 1(b) shows the simulation results of the three transmission schemes (compounding RC, compounding CR, and compounding RC+CR) at the volume rate of 400 Hz as listed in Table I. Due to the symmetrical layout of the row elements and column elements, the PSF of the RC scheme is identical to that of the CR scheme except the image is rotated by 90°. Because the compounding RC+CR scheme is a combination of the RC scheme and CR scheme, the PSF is symmetrical in the x-y plane. Since the RC+CR scheme used a smaller angular aperture (for x-z or y-z plane) as compared to RC or CR to maintain the same volume-rate, it had weaker transmit focusing and a wider main lobe (i.e., worse spatial resolution). The average Full Width at Half Maximum (FWHM) along y-direction was calculated using 30 point targets located at depths between 20 – 50 mm. Figure 1(c) shows the similar simulation results of the three transmission schemes at the volume rate of 2000 Hz. Due to the limited number of compounding angles (e.g., 7 angles), the main lobe along the transmit focusing direction was wide. The compounding RC+CR scheme had a better B-mode spatial resolution (measured at -6 dB) in both x- and y-direction than either RC or CR alone.

The same 3D SWE detection sequences were then used to image a multi-purpose multi-tissue phantom, as shown in Figure 2(a) and (b). The FWHM was measured for the wire target located at 41 mm depth and averaged along the x-direction. Similar to the results obtained from the simulation, the RC scheme or the CR scheme had better B-mode resolution than the compounding RC+CR scheme at 400 Hz, while the compounding RC+CR scheme had better resolution at 2000 Hz.

**Figure 2:**
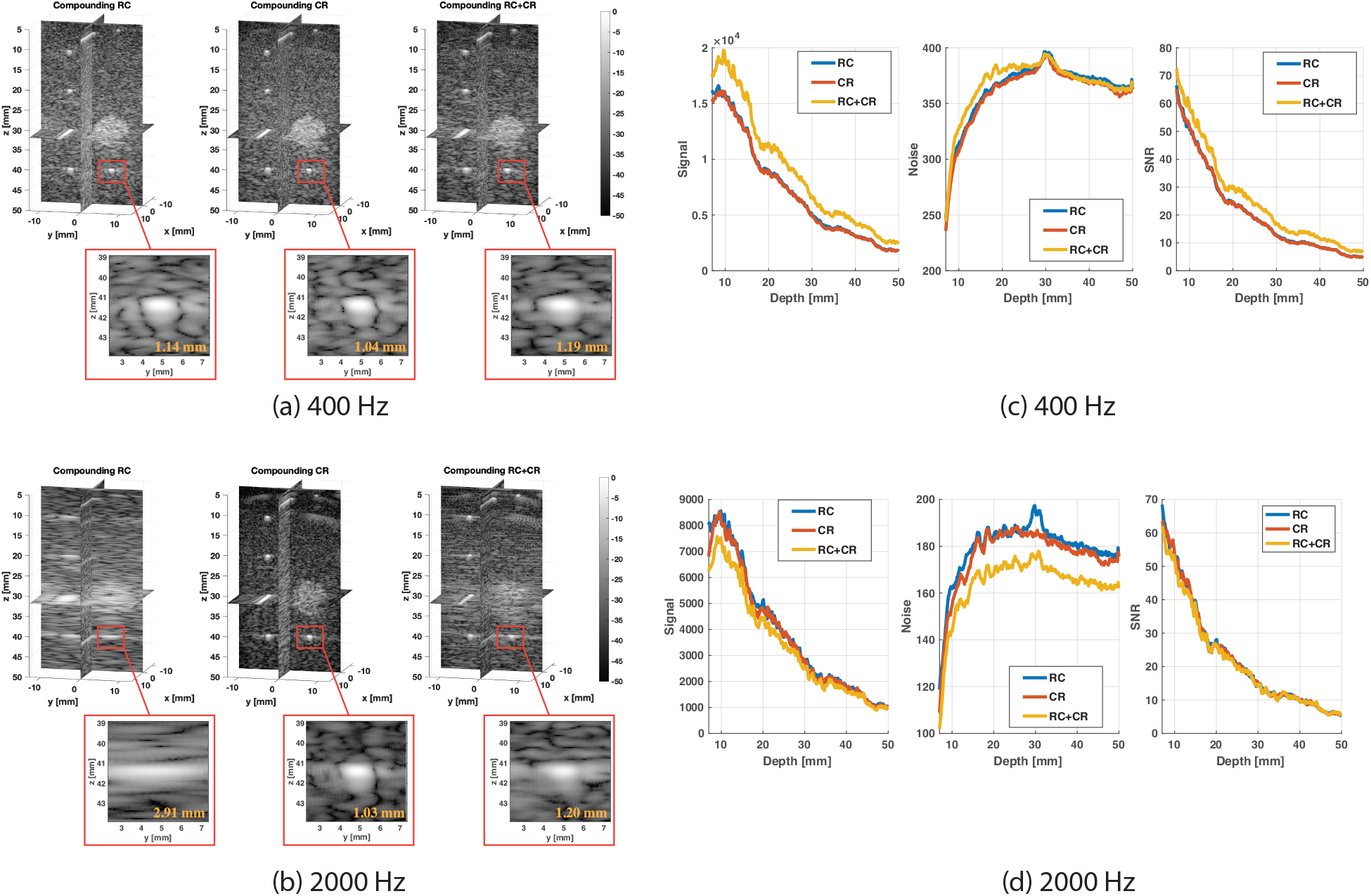
Volumetric images and experimental SNR of the phantom. (a) Volumetric images (slice view) of the multi-purpose multi-tissue phantom using different transmission schemes (compounding RC, compounding CR, and compounding RC+CR) at 400 Hz, as listed in Table I. Magnified view (y-z slice) of the PSF at 41 mm depth is presented with measured FWHM along the y-direction, and the FWHM is averaged along the x-direction. (b) Volumetric images of the phantom using different transmission schemes at 2000 Hz, as listed in Table I. (c) Signal, noise, and SNR measurements in the multi-purpose phantom using different transmission schemes at 400 Hz, as listed in Table I. The measurement was conducted on a region without hyperechoic objects. (d) Signal, noise, and SNR measurements in the phantom using different transmission schemes at 2000 Hz.

Another important metric for 3D SWE is the signal-to-noise ratio (SNR) for shear wave detection, which affects the quality of the detected shear wave signal, especially when the motion is weak such as in the case of ARF-induced shear waves. Figure 2(c) and (d) show the measured signals, noises, and SNRs of the three sequences as a function of imaging depth. Due to the symmetry of RC and CR schemes, the SNR performances were identical for these two schemes. Because RC+CR scheme used a smaller angular aperture that distributes more acoustic energy within the field of view (FOV), SNR was higher for RC+CR than either RC or CR when the volume rate was 400 Hz (see Figure 2(c)).

### External Vibration-based 3D SWE

After studying the RCA-based detection sequence for 3D SWE, we then performed the 3D shear wave study using a homogenous phantom and an elasticity phantom (see ‘External Vibration-based 3D SWE’ in Materials and Methods). The imaging sequence details are reported in for each application. Figure 3(a) shows the detected 3D shear wave motion at 100 Hz in the homogenous phantom (i.e., the multi-purpose multi-tissue phantom) with a volumetric imaging rate of 400 Hz. Figure 3(b) shows the reconstructed shear wave speed (SWS) volume using the 3D local frequency estimation (LFE) method, and Figure 3(c) shows the corresponding histogram of SWS from the full volume. The average SWS estimated using the proposed 3D SWE method was 2.38 ± 0.35 m/s, while the SWS measured by the LOGIQ E10 ultrasound system (General Electric Healthcare, Wauwatosa, WI, USA) was 2.98 m/s. The nominal SWS of the phantom according to the literature [16] is 2.60 ± 0.15 m/s. The measurement using the GE system was conducted with an L2-9-D linear array (9 MHz; General Electric Healthcare, Wauwatosa, WI, USA) and the ARF-based 2D elastography mode [28]. The underestimation of SWS using the proposed method may be explained by the lower shear wave frequency (i.e., 100 Hz) from using the external vibrator.

**Figure 3:**
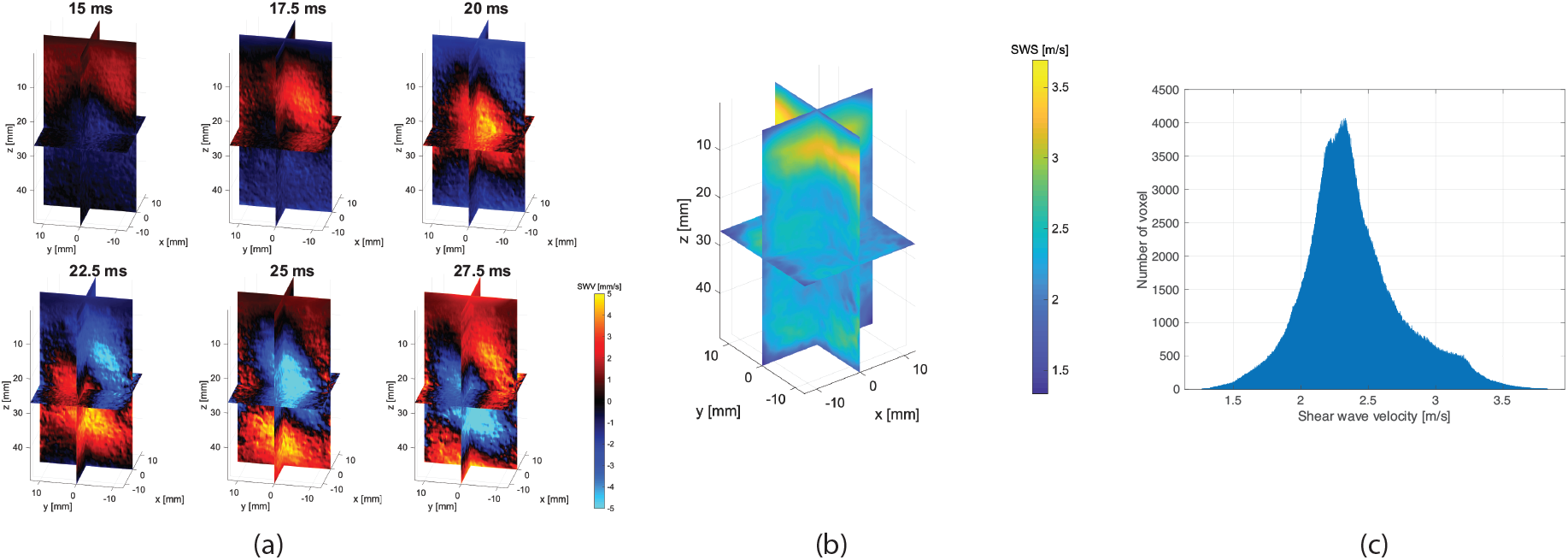
Shear wave motion and reconstructed SWS map of the homogenous phantom using the external vibration-based 3D SWE. (a) Volumetric images (slice view) of the propagating shear waves induced by external vibration in the homogenous phantom at different time points. The 100 Hz continuous shear wave was detected by the RCA array with a volume rate of 400 Hz. (b) Reconstructed SWS map (slice view) of the homogenous phantom. (c) Histogram of the full volume of SWS values.

For the elasticity phantom study, a stiff spherical lesion with Young’s modulus of 64.9 kPa (4.65 m/s) was imaged, and Young’s modulus of the background is 18 kPa (2.45 m/s). Figure 4(a) shows representative volumetric frames of detected shear wave motion (200 Hz) as a function of time. Figure 4(b) shows the reconstructed SWS volume of the elasticity phantom with clear visualization of the spherical lesion, the artifact located at the left top corner is probably due to a boundary effect. The measured average SWS of the background was 2.67 m/s with the GE system (Figure. S1) and 2.43 ± 0.19 m/s with the proposed method; the SWS of the stiff target was 4.94 m/s with the GE system (Figure. S1), and 3.30 ± 0.06 m/s with the proposed 3D-SWE method.

**Figure 4:**
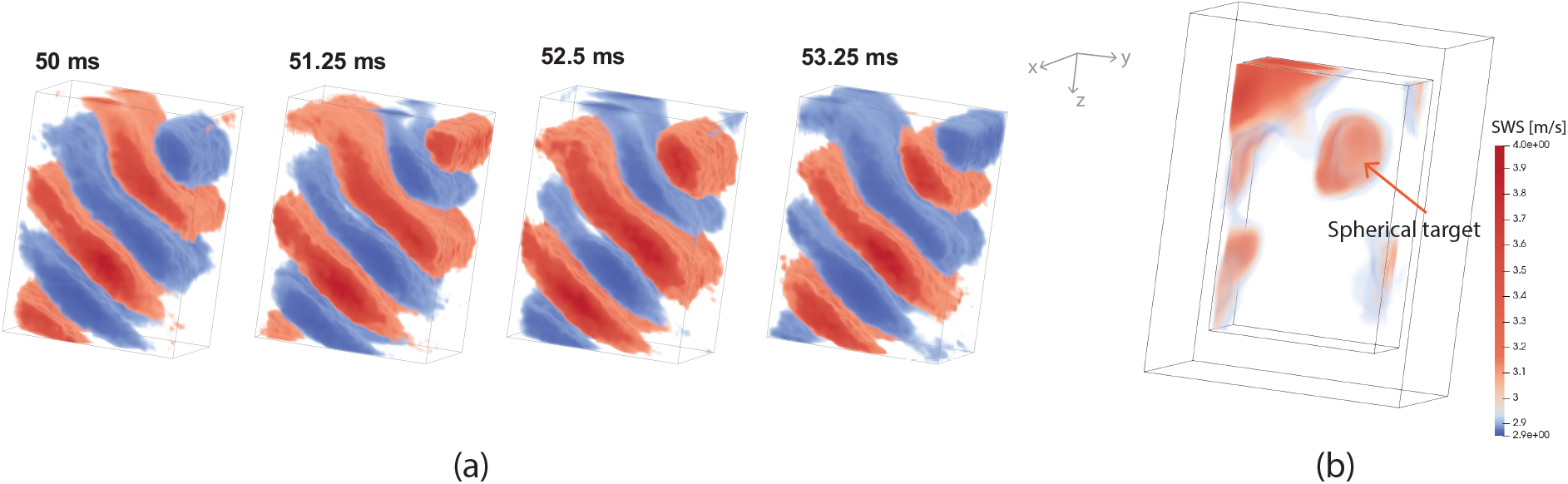
Shear wave motion and SWS map of the elasticity phantom using the external vibration-based 3D SWE. (a) Representative volumetric frames (25.6 × 25.6 × 40 mm^3^) of shear wave motion using external vibration in the elasticity phantom at four consecutive time points. The 200 Hz continuous shear wave was detected by the RCA array with a volume rate of 800 Hz. (b) Volumetric image of reconstructed SWS map of the elasticity phantom. The artifact located at the left top corner is probably due to a boundary effect.

### ARF-based 3D SWE

Finally, we applied the proposed method to detect the ARF-based shear wave signal. Because of the relatively low element sensitivity of the RCA array used in this study (see Figure. S2), it was challenging to use the same array to generate shear waves. This is only a limitation with the particular RCA array used in this study and does not necessarily apply to other RCA probes. Due to this limitation, we used a 1D linear array transducer (L7-4, Philips Healthcare, Andover, MA, USA) to produce the push beam for shear wave generation for the RCA to detect. Figure 5(a) shows the detected shear wave motion in the x-y slice at 1.5 ms using compounding RC and compounding RC+CR scheme, as listed in Table II. The detected shear wave using compounding RC scheme has better signal quality compared to the RC+CR scheme, although the RC+CR scheme achieves better B-mode spatial resolution at 2000 Hz (see Figure 2(b)). Figure 5(c) shows the detected shear wave motion at six consecutive time points using the compounding RC scheme at a volume rate of 2000 Hz. The ARF-induced cylindrical-shaped shear wave by the L7-4 linear array can be clearly visualized. The estimated SWS of the homogenous phantom using the random sample consensus (RANSAC) method was 3.05 m/s, as shown in Figure 5(b). The SWS measured using the GE system was 2.75 m/s.

**Table II.** Imaging specifications of the external vibration-based 3D SWE and ARF-based 3D SWE studies

| Specs | External Vibration-based 3D |  | ARF-based 3D SWE |  |
| --- | --- | --- | --- | --- |
|  | Multi-purpose phantom | Elasticity phantom |  |  |
| Transmission scheme | Compounding RC | Compounding RC | Compounding RC | Compounding RC + CR |
| Number of transmissions | 31 | 17 | 7 | 6 |
| Angular pitch [°] | 1 | 1 | 1 | 1 |
| Maximum angular aperture [°] | 15 | 8 | 3 | 1 |
| Pulse repetition frequency, PRF [kHz] | 12.8 | 13.6 | 14 | 12 |
| Volume rate [Hz] | 400 | 800 | 2000 | 2000 |
| External vibration frequency | 100 | 200 | / | / |
| Number of volumes acquired | 40 | 80 | 20 | 20 |
| Acquisition time [ms] | 100 | 100 | 10 | 10 |

**Figure 5:**
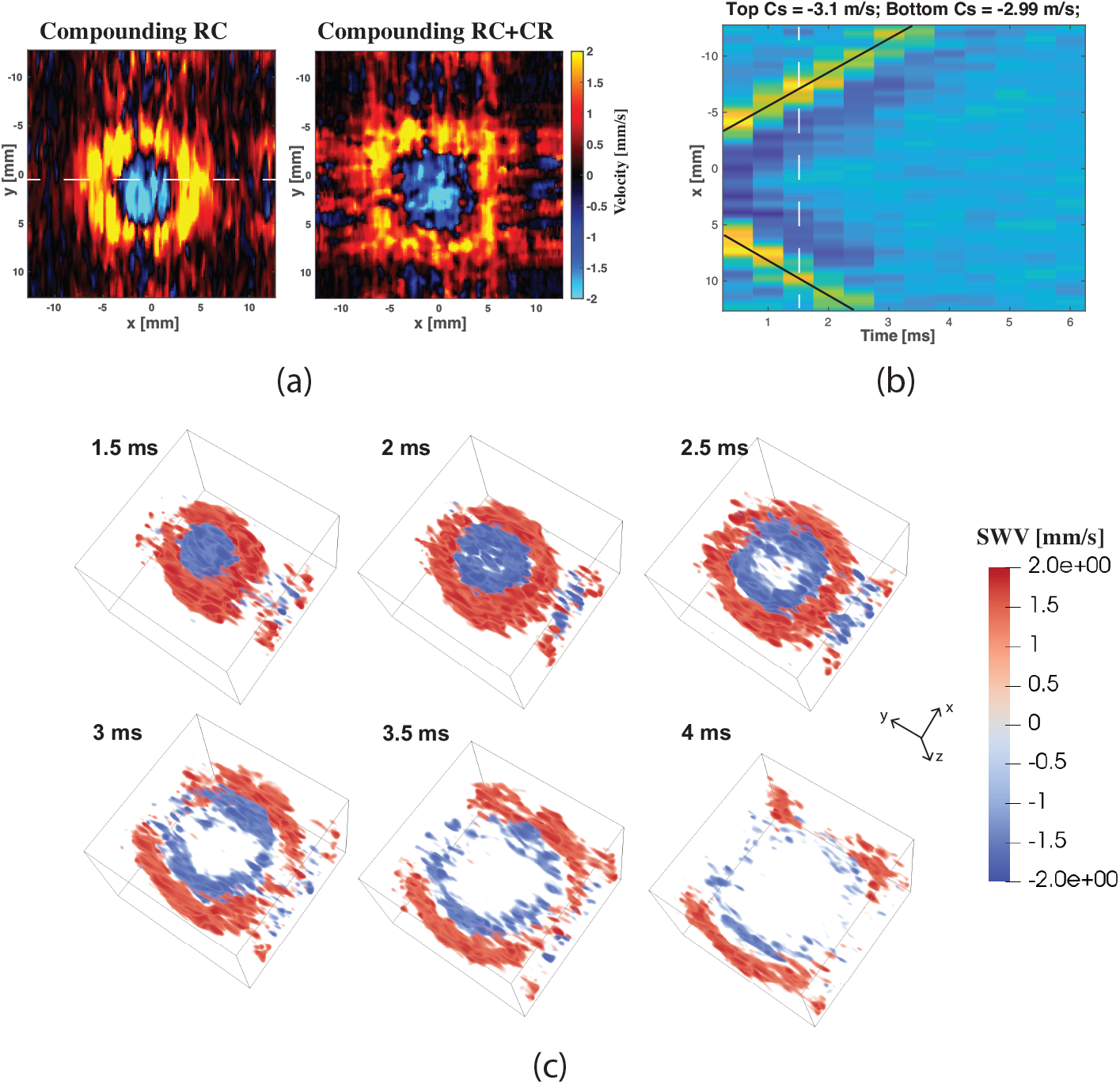
Shear wave motion using the ARF-based 3D SWE. (a) Detected ARF-based shear wave motion (x-y slice) at 1.5 ms using the compounding RC and compounding RC+CR scheme as listed in Table II. (b) 1-D shear wave motion induced by ARF along the x-direction over time, and the SWS speed was estimated using the random sample consensus (RANSAC). (c) Representative volumetric frames (25.6 × 25.6 × 14.8 mm^3^) of shear wave motion at six consecutive time points.

## Discussion

Currently, 3D SWE is challenged by various technical limitations such as the limited volume rate of 3D imaging systems and the high fabrication and computation costs associated with conventional 2D matrix arrays. These limitations remain the major challenge for 3D elastography in clinical practice. In this article, we introduced a 3D-SWE method based on the RCA array with shear waves induced by either external vibration or ARF. The phantom studies demonstrate that the proposed method is capable of robust 3D shear wave tracking at high volume rates (e.g., 2000 Hz) without the need for repeated shear wave generation and detection.

The main advantage of RCA-based 3D SWE is that the RCA array is compatible with most commercial ultrasound systems because of its lower channel count, and the RCA array has lower fabrication costs compared to conventional 2D matrix arrays [26]. For example, our RCA array includes 128 + 128 elements with a 25.4 × 25.4 mm^2^ footprint, while for a 2D matrix array 128 × 128 elements (i.e., 16,384 channels) would be needed for the same footprint. Meanwhile, a distinct advantage of the RCA array is that it is feasible to achieve a high imaging volume rate (e.g., 1000 – 2000 Hz) that is challenging for conventional 2D matrix arrays to achieve. Moreover, because 2D RCA arrays do not require multiplexing, they can better sustain the long push pulses for ARF-based SWE. In theory, there should be no technical barriers for RCA to transmit long-duration push pulses.

In this study, three different transmission schemes (i.e., compounding RC, compounding CR, and compounding RC+CR) were evaluated *in silico* and *in vitro*. We found that with a fixed angular pitch (i.e., 1°) and a relatively low volume rate (e.g., 400 Hz), the compounding RC scheme or compounding CR scheme achieved better B-mode spatial resolution as compared to the compounding RC+CR scheme, which uses a smaller angular aperture due to the limited number of transmissions (Figure 2(a)). Meanwhile, higher energy gain was achieved by the RC+CR scheme (Figure 2(c)) because of smaller angular aperture, which showed better shear wave detection SNR. In this study, the compounding RC scheme was used for shear wave detection with external vibration due to the better B-mode resolution, as listed in Table II, which is preferred for detecting high-quality shear waves to reconstruct small objects like the spherical lesion. However, the resolution difference between RC and RC+CR was not significant (Figure 1(b)). For the ARF-based SWE, both compounding RC and compounding RC+CR scheme were investigated, and the detection using compounding RC achieved better shear wave signal quality (Figure 5(a)).

The tradeoff between spatial resolution and SNR needs to be balanced for different applications associated with different volume rates. For example, to detect a small lesion, shear waves with small wavelengths are necessary, and a sequence with high B-mode resolution is preferred because of the more robust spatial sampling. However, if the SNR is more important than spatial resolution (e.g., weak shear wave signal from deep tissues) then a sequence with higher SNR is preferred. For example, one can use the RC+CR scheme or the RC scheme with a reduced angular aperture to distribute maximum acoustic energy within the FOV. Alternatively, the number of transmissions (i.e., number of compounding angles) can be increased, and multiple acquisitions with repeated pushes or vibrations can be used to further boost shear wave SNR. Other advanced transmission schemes can also be applied to improve SNR such as the Hadamard-encoding method [29].

Since RCA is only capable of one-way focusing, the imaging quality is always compromised when the number of compounding angles is reduced to allow for higher imaging volume rate. In our study, a 2000 Hz volume rate (with 6 or 7 compounding angles) was used to detect the ARF-induced shear waves, which lead to noisy shear wave signals (Figure 5) that were challenging for SWS estimation methods such as LFE and cross-correlation. To address this issue, multiple acquisitions with time averaging can be used to improve the shear SNR, although in practice for human imaging a breath hold may be necessary to avoid motion artifacts. Recently several RCA-based adaptive beamforming methods were proposed to improve the RCA imaging quality by taking advantage of the RCA array layout [30, 31]. However, these methods generally involve multiplication between the RC and CR data, which distorts the phase information and is not suitable for detecting shear wave motion.

Apart from compounding plane wave imaging, other transmission schemes can also be used on the 2D RCA array for 3D SWE. For example, synthetic aperture imaging [17] can offer optimal spatial resolution (see Figure. S3 (d)-(f)) and line-by-line focused beam imaging (see Figure. S3 (g) and (h)) can offer optimal SNR performance. Wide beam imaging is another option that offers both high shear wave detection SNR and spatial resolution (see Figure. S3 (i) and (j)). However, one major limitation of these three methods is that a large number of transmissions is needed, which results in a volume rate that may be too low to be used for 3D SWE. Nevertheless, these sequences may be useful for external vibration-based SWE where shear wave frequency is low. Alternatively one can use repeated cycles of shear wave generation and detection to synthesize shear wave data with these low volume rate sequences [28].

In this study, for the first time to the best of our knowledge, RCA-based 3D SWE with external vibration or ARF was developed and verified on phantoms with the clinic-ready ultrasound imaging system, and the quantitative measurements were compared with a state-of-the-art clinical 2D ultrasound SWE system. The proposed method can provide 3D elastography at a high volume-rate (e.g., 2000 Hz) with short acquisition time (e.g., tens of milliseconds) and fast processing speed by GPU-based computation acceleration. Ongoing studies are being conducted to use a new probe for ARF-based 3D SWE and evaluate its performance for *in vivo* imaging.

## Materials and Methods

### Imaging sequence for 3D shear wave detection

To detect the shear waves generated by either ARF or external vibration, a high volume-rate imaging with adequate imaging quality is necessary. The RCA array includes two groups of orthogonally arranged 1-D linear arrays (i.e., rows and columns), which transmit and receive in an alternate fashion for 3D imaging (e.g., rows transmit, and columns receive, and vice versa). One limitation of such an arrangement is that RCA arrays only support one-way focusing. Figure 1(a) shows an example of transmission of steered plane waves in the y-z plane (using the row elements) and receive of the plane waves in the x-z plane (with the column elements). In this case, only transmit focusing can be achieved in the y-z plane (i.e. there is no y-z receive focusing), and only receive focusing can be achieved in the x-z plane (i.e. there is no x-z transmit focusing).

Among various transmission schemes, such as compounding plane-wave imaging [14, 32], synthetic aperture imaging [17], and line-by-line focused beam imaging, compounding plane-wave imaging provides the best tradeoff for high volume-rate scanning because it presents a balanced performance between imaging quality and imaging frame-rate. However, due to the limitation of one-way focusing, the point spread function (PSF) of either scheme is asymmetric (e.g., elongated along the y-direction when using the RC scheme). A common approach to improve the spatial resolution and obtain a symmetric PSF is to use the RC + CR scheme. However, this method reduces the imaging volume-rate by half because it requires twice as many pulse-echo data acquisition cycles. To maintain the same volume-rate as the RC or CR scheme, the RC+CR scheme uses half the number of compounding angles in either x-z or y-z plane, and the angular aperture will be reduced by half with a fixed angular pitch.

To study the impact of compounding RC, compounding CR, and compounding RC+CR schemes for 3D SWE (e.g., spatial resolution, SNR), three corresponding sequences with the same volume rate of 400 Hz or 2000 Hz and a fixed angular pitch of 1° were designed and tested to identify the suitable transmission method for robust 3D shear wave detection and tracking. The 400 Hz volume rate is more appropriate for shear wave detection with external vibration, and the 2000 Hz volume is appropriate for the ARF-based shear wave detection. Table I summarizes the configurations of the studied sequences. Both simulation and phantom studies were carried out to evaluate the imaging performance of each sequence. Simulations were conducted using the Verasonics simulator with multiple point targets located at different depths (e.g., 20 – 50 mm). The simulation settings were specified to match the transducer parameters which were used for the phantom study. For the phantom study, a Verasonics Vantage 256 system (Verasonics Inc., Kirkland, WA, USA) and a 128 + 128 element 2D RCA array transducer (Vermon SA, Tours, France; central frequency at 5.2 MHz (measured at -20 dB)) were used for the acquisitions. The transmit frequency was 5.2 MHz with a sampling frequency of 20.8 MHz. A multi-purpose multi-tissue ultrasound phantom (Model 040GSE, CIRS Inc. Norfolk, VA, USA) was used for the resolution and SNR evaluation.

The FWHM of the point target was used to evaluate the spatial resolution. To quantify the experimental SNR performance (i.e., for shear wave detection), 100 volumes were acquired from a homogeneous phantom and a region of the phantom with uniform speckle patterns were used for SNR estimation. The image noise *n*(*x*, *y*, *z*) is calculated as the time standard deviation of the volumetric image *I*(*x*, *y*, *z*, *t_i_*)

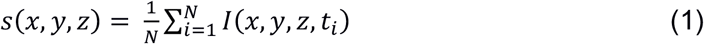

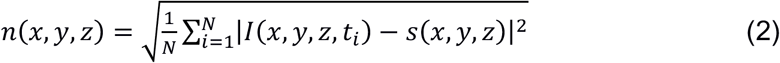

where *s*(*x*, *y*, *z*) is the mean signal over time, and *N* is the number of volumes. The signal, noise, and SNR were averaged along the x- and y-direction to obtain an accurate profile of the SNR as a function of the depth [33], and the SNR is given by

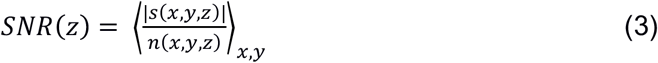

Based on the quantitative evaluation of the resolution and SNR *in silico* and *in vitro*, proper transmission schemes were selected and used for the rest of the studies.

### External Vibration-based 3D SWE

An electromechanical vibrator (Bruel & Kjaer VTS Ltd., Royston, UK) was used to generate harmonic shear waves with different frequencies (e.g., 100 Hz – 200 Hz) from a function generator (33210A, Keysight Technologies Inc., Santa Rosa, CA, USA). The experiment setup is illustrated in Figure 6(a). The sinusoidal signal from the function generator was amplified by an amplifier (XLS 2502, Crown Audio, Los Angeles, CA) before being sent to the vibrator. A metal rod with a bar-shaped tip was used to conduct the vibration from the vibrator to the phantom. The vibrator was synchronized with the ultrasound system. Both the multi-purpose phantom (040GSE) and an elasticity phantom (Model 049, CIRS Inc. Norfolk, VA, USA) were utilized to test the 3D-SWE imaging performance.

**Figure 6:**
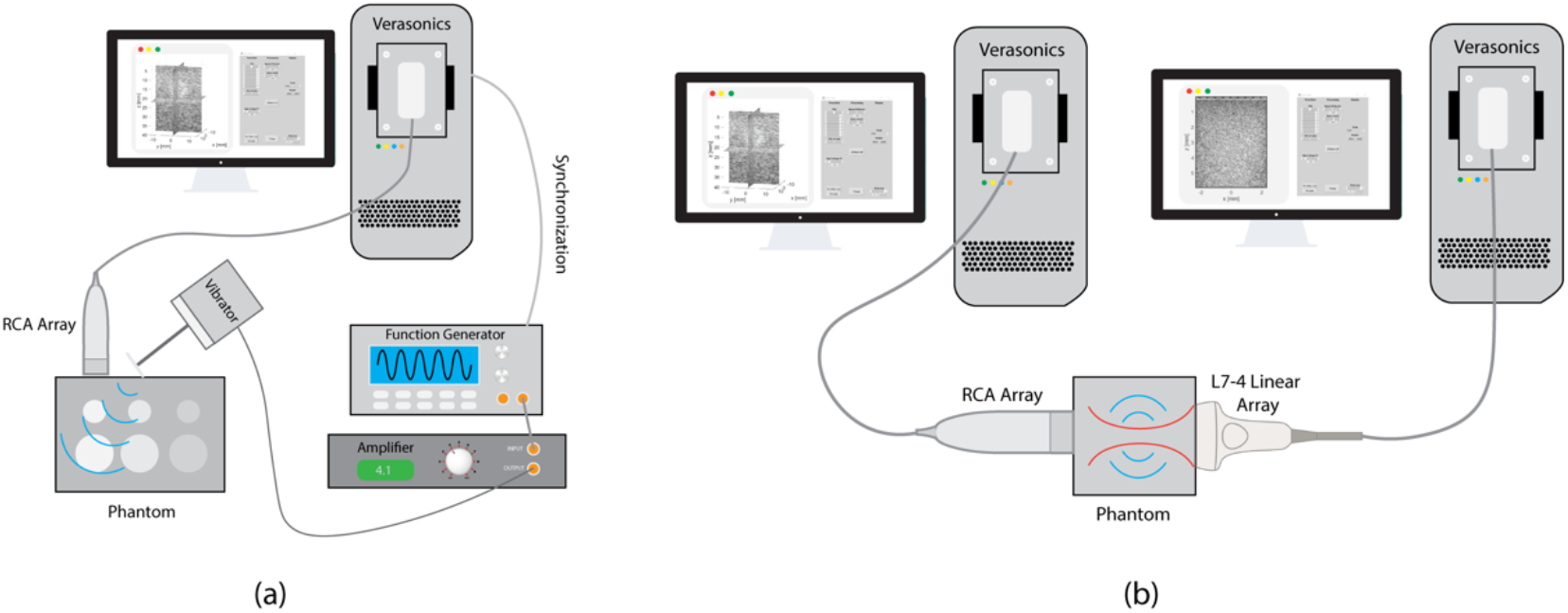
Experimental setups. (a) Experimental setup of the external vibration-based 3D SWE study. A vibrator with a bar-shaped tip was fixed and driven by amplified sinusoidal signals to generate continuous shear waves delivered into the phantom tissue, and the synchronized RCA array performed volumetric imaging during the vibration. (b) Experimental setup of the ARF-based 3D SWE study. The L7-4 linear array generated a shear wave by acoustic radiation force into the homogenous phantom, and the synchronized RCA array performed volumetric imaging. Two Verasonics Vantage systems were used to drive the L7-4 and RCA arrays. The L7-4 linear array was aligned along the x-direction of the coordinates defined by the RCA array.

To avoid aliasing, the volume-rate of the detection sequence was set to four times the shear wave frequency, as listed in Table II. For the homogenous phantom study (i.e., the multi-purpose multi-tissue phantom), a 100 Hz shear wave was generated and detected with a scanning rate of 400 Hz. For the heterogeneous phantom study (i.e., the elasticity phantom), a stiff spherical lesion (10 mm diameter) was imaged. Because the resolution of the shear wave imaging is fundamentally limited by the wavelength of the shear wave, to resolve the spherical lesion from the background with better resolution, a 200 Hz shear wave was used and detected by an 800 Hz RCA detection sequence.

In this study, we developed a GPU-based beamforming method for fast ultrasound data reconstruction. In-phase/quadrature (IQ) data were beamformed and then used to calculate the particle velocity induced by shear waves using the autocorrelation method [34]. To alleviate shear wave interference inside the phantom from external vibration, a 3D directional filter was applied on the shear wave data to extract shear wave signals that propagate away from the source [35]. 3D SWS maps were reconstructed using the 3D local frequency estimation (LFE) method [36, 37] or 3D local cross-correlation methods [19].

### ARF-based 3D SWE

The experiment setup is depicted in Figure 6(b). The two transducers were carefully aligned and operated by two Verasonics Vantage systems, which were synchronized. A homemade homogenous tissue-mimicking phantom was sandwiched between the two transducers. The L7-4 transmitted the push beam (push beam central frequency, 4 MHz; F-number, 1; focal depth, 3 cm; push length, 600 us) immediately after the RCA started data acquisition. Similarly, 1-D autocorrelation was used for shear wave motion calculation. Due to the low sensitivity of the RCA array (see Figure. S2) and the high volume-rate used for detection (i.e., 2000 Hz), the shear wave signal was noisy due to low SNR from the limited number of transmissions (e.g., 7 transmissions), especially along with the transmit focusing direction (e.g., y-direction) when only the RC scheme was applied. Furthermore, the ARF-based shear wave motion is much weaker than the external vibration. To best estimate the SWS of the phantom from noisy shear wave signal, the RANSAC [38] method was used to fit the shear wave motion and calculate SWS.

## Author contributions

Z.D., S.C., and P.S. generated the idea and designed the experiments. Z.D., H.C., S.C., and P.C. developed and applied the shear wave motion detection and shear wave speed estimation algorithms. Z.D., J.K., and M. R. L. performed the experiments. All authors were involved in the data analysis, critical revision of the manuscript, and discussion.

## Data availability

All data are available within the Supplementary Materials, or available from the authors upon request.

## Supplementary Materials

**Figure. S1.**
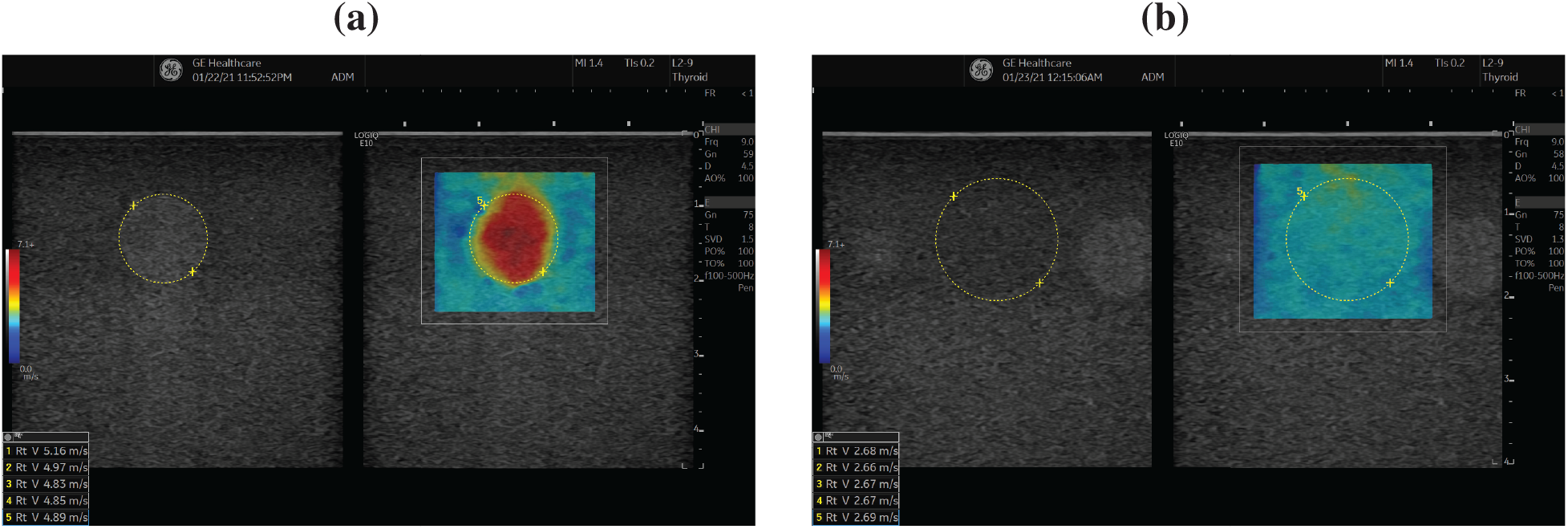
Shear wave speed maps of the (a) stiffer spherical lesion and (b) background of the elasticity phantom using the GE LOGIC E10 system and a 1D linear array. The measurement was conducted in ARF-based 2D SWE mode.

**Figure. S2.**
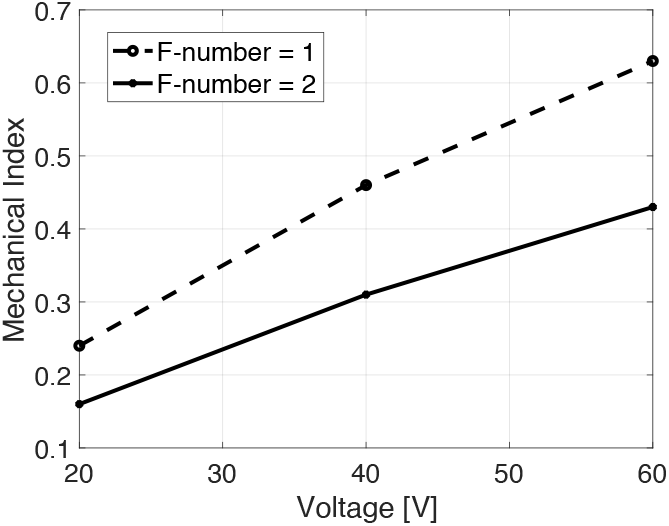
Mechanical index of the RCA array as a function of the input voltage. The measurements were performed using a needle hydrophone in the water tank, and the RCA transmitted a focused beam using 64 elements (F-number = 2) or 128 elements (F-number = 1), and the focal depth is 25 mm.

**Figure. S3.**
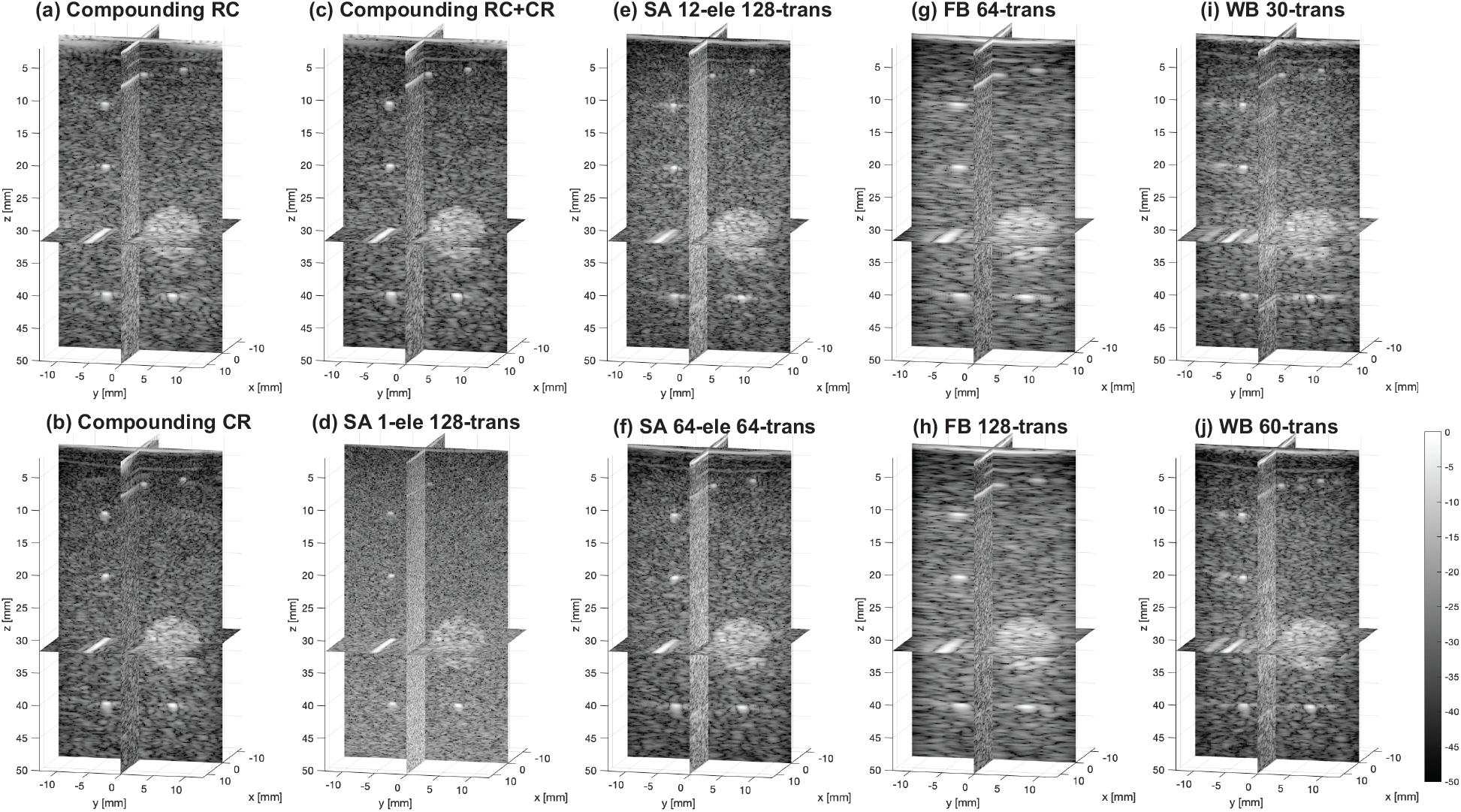
Reconstructed volumetric images (slice view) of the phantom using four different transmission schemes (compounding plane wave imaging, synthetic aperture imaging (SA), focused beam imaging (FB), and wide beam imaging (WB)). (a) Compounding plane wave imaging with RC scheme with an angular pitch of 1°, angular aperture of 15°, 31 compounding angles, and 400 Hz volume rate. (b) Compounding plane wave imaging with CR scheme with an angular pitch of 1°, angular aperture of 15°, 31 compounding angles, and 400 Hz volume rate. (c) Compounding plane wave imaging with RC+CR scheme with an angular pitch of 1°, angular aperture of 15°, 62 compounding angles, and 200 Hz volume rate. (d) SA imaging with 128 transmissions and 100 Hz volume rate, one element was used for each transmission. (e) SA imaging with 128 transmissions and 100 Hz volume rate, virtual source (a sub-aperture of 12 elements) was used for each transmission. (f) SA imaging with 64 transmissions and 200 Hz volume rate, virtual source (a sub-aperture of 64 elements) was used for each transmission. (g) FB imaging with 64 transmissions and 200 Hz volume rate, 32 elements were used for each focused beam with a F-number of 4. (h) FB imaging with 128 transmissions and 100 Hz volume rate, 32 elements were used for each focused beam with a F-number of 4. (i) WB imaging with 30 transmissions and 400 Hz volume rate, 50 elements were used for each sub-region compounding. (j) WB imaging with 60 transmissions and 200 Hz volume rate, 50 elements were used for each sub-region compounding.

